# Synthesis, secretion, and perception of abscisic acid regulates stress responses in *Chlorella sorokiniana*

**DOI:** 10.1101/180547

**Authors:** Maya Khasin, Rebecca E. Cahoon, Sophie Alvarez, Richard Beckeris, Seong-il Eyun, Qidong Jia, Jean-Jack Riethoven, Kenneth W. Nickerson, Wayne R. Riekhof

## Abstract

Abscisic acid (ABA) is a phytohormone that has been extensively characterized in higher plants for its roles in seed and bud dormancy, leaf abscission, and stress responses. Genomic studies have identified orthologs for ABA-related genes throughout the *Viridiplantae*, including in unicellular algae; however, the role of ABA in algal physiology has not been characterized, and the existence of such a role has been a matter of dispute. In this study, we demonstrate that ABA is involved in regulating algal stress responses. *Chlorella sorokiniana* strain UTEX 1230 contains genes orthologous to those of higher plants which are essential for ABA biosynthesis, sensing, and degradation. RNAseq-based transcriptomic studies reveal that treatment with ABA induces dramatic changes in gene expression profiles, including the induction of a subset of genes involved in DNA replication and repair, a phenomenon which has been demonstrated in higher plants. Pretreatment of *C. sorokiniana* cultures with ABA exerts a protective effect on cell viability in response to ultraviolet radiation. Additionally, *C. sorokiniana* produces and secretes biologically relevant amounts of both ABA and the oxylipin 12-oxo-phytodienoic acid (OPDA) into the growth medium in response to abiotic stressors. Taken together, these phenomena suggest that ABA signaling evolved as an intercellular stress response signaling molecule in eukaryotic microalgae prior to the evolution of multicellularity and colonization of land.

## Introduction

Plants respond to abiotic stresses in multiple and nuanced ways, involving carefully regulated and interconnected signaling pathways. In particular, the response to saline and drought stress – stressors characterized by lowered water availability and desiccation– is mediated by a number of small-molecule signals and many interacting signal transduction pathways. These signals and pathways include reactive oxygen species (ROS), salt overly sensitive genes (SOS), 12-oxophytodienoic acid (OPDA), jasmonic acid (JA), and abscisic acid (ABA) [1]. ABA is a molecule found throughout all domains of life except Archaea [2]. In plants, it functions as a stress signaling molecule originally discovered and named for its role in leaf abscission, which is integral to their response to many types of abiotic stresses related to water availability, including drought/desiccation, salinity, and cold stress [3]. The SOS pathway, which contributes to the salinity response, relies upon calcium sensing, of which ABA is an agonist. Briefly, SOS3 is a Ca^2+^ sensor which activates SOS2, a Ser-Thr kinase, which leads to the recruitment of SOS1, a Na^+^/H^+^ antiporter. ABA production is additionally induced by singlet oxygen stress as a byproduct of photosynthesis, and further transduces ROS signaling by inducing H_2_O_2_, which mediates stomatal closure in plants.

Orthologs to genes involved in plant hormone signaling pathways, including ABA signaling, have been reported in algal genomes, but at present little functional information is available about the role of ABA in algae. Orthologs to genes implicated in ABA signaling have been observed in *Chlorella variabilis* NC64A[4], the streptophyte *Klebsormidium flaccidum*[5], the picoeukaryotic marine alga *Ostreococcus tauri*[6], and the model chlorophyte *Chlamydomonas reinhardtii* (2, 7). Though putative plant hormone related orthologs, including ABA related genes, have been described in algae [2], the specific functions of ABA and its physiological and evolutionary significance have remained unclear. ABA itself has been identified in extracts of algae from a number of eukaryotic lineages, and it has also been identified in cyanobacteria, some of which induce the formation of nitrogen-fixing heterocysts in response to exogenous ABA (8, 9). *Dunaliella* species have been demonstrated to accumulate ABA and secrete it into the growth medium upon saline or alkaline shock [8]. These experiments implicated ABA signaling in β-carotene accumulation and salt stress tolerance [10], leading to the proposition that microalgal ABA signaling may share some functions with this pathway in higher plants.

In higher plants, ABA signaling is mediated by PYR1/PYL/RCAR [11], a series of soluble ABA-binding proteins that inactivate SNF1-related protein kinases (SnRK2s). Additional ABA perception functions are mediated by chloroplast localized magnesium chelatase subunit H (ChlH), a member of the porphyrin biosynthesis pathway with a role in plastid to nucleus retrograde signaling [12], and GTG1, a GPCR-like transmembrane G protein [13].

ABA signaling in higher plants also integrates with other stress-related signaling pathways, including reactive oxygen species (ROS), salt overly sensitive (SOS) signaling, as well as the lipid-derived phytohormones 12-oxo-phytodienoic acid (OPDA) and jasomonic acid (JA) [14]. Specifically, the saline stress response in higher plants involves SOS and ROS signaling, both of which interact with each other and with ABA mediated pathways (1,15). Transcriptomic analyses in *Arabidopsis* have revealed ABA dependent and independent gene expression in response to salt and drought stress [16]; additionally, genotoxic stress, DNA damage repair, and ROS signaling may have ABA independent components (17, 18). There are ABA dependent and independent features of these signaling pathways in *Arabidopsis* [19]. In Arabidopsis, both salinity and genotoxic stress signaling pathways are interconnected by MAP kinases. *C. sorokiniana* encodes putative orthologs for a complete SOS pathway as well as orthologs for interacting members of remaining stress response pathways, suggesting an ancient evolutionary origin of plant stress response pathways. In Arabidopsis, MAP kinase phosphatase 1 (MKP1) is implicated in both salt and genotoxic stress signaling pathways [20]. Mutant *mkp1* plants are still resistant to salt stress but more highly sensitive to genotoxic stress.

In this study, we demonstrate that *C. sorokiniana* secretes physiologically relevant concentrations of ABA and OPDA during osmotic stress; furthermore, 290 transcripts are differentially expressed in response to treatment with ABA, and a large portion of the upregulated genes are involved in DNA replication and repair. However, in contrast with higher plants, ABA treatment induces no loss in cell viability, growth rate, or cell yield. Pretreatment of cultures with ABA was found to increase tolerance of cultures to UV irradiation, but not consistently to salt stress. We further present comparative genomic evidence for the presence and function of ABA and related signaling genes in *C. sorokiniana*. Our work thus integrates multiple approaches to demonstrate that a plant hormone acts an intercellular signaling molecule in unicellular eukaryotic algae. More specifically, we propose that the role of ABA as a stress response phytohormone predates the evolution of multicellularity and may have had a pivotal role in the colonization of land.

## Results

### *Chlorella sorokiniana* encodes orthologs to phytohormone-mediated stress signaling pathways

The *Chlorella sorokiniana* strain UTEX-1230 was identified as a promising biomass and waste-remediation organism in a large scale screening study conducted by a consortium of algal biologists at the University of Nebraska [21–23]. Its genome was subsequently sequenced, and a combination of *in silico* gene-prediction programs and RNA sequencing was used to annotate predicted genes, open reading frames, and protein sequences. The sequencing and annotation of this strain will be described in detail in a subsequent publication, (Cerutti et al., in preparation); however, a current gene catalog including the 11 protein sequences used in our analysis (Tables 1 and 2) has been deposited in Genbank.

To identify and catalog phytohormone related genes, reciprocal BLAST searches were performed using experimentally verified protein sequences from seed plants to search the predicted *C. sorokiniana* proteome. We initially analyzed and cataloged genes related to auxin, cytokinin, ethylene, JA, and ABA signaling, and in this study we focus on genes mediating the synthesis and signaling functions of ABA and 12-OPDA, a bioactive precursor of JA [24].

*C. sorokiniana* genes related to ABA biosynthesis, transport, sensing, and degradation are listed in Table 1, along with their predicted functions. The ABA related genes we identified are closely related to genes for ABA synthesis and signaling in higher plants, although the canonical seed-plant ABA receptor family (PYR/PYL/RCAR) along with downstream signaling components were absent. Phylogenetic analyses (Figs. 1-5) demonstrate that these genes constitute lineages that are more closely related to plant ABA signaling genes than they are to paralogous genes containing similar domains. Though no PYR1/PYL/RCAR like proteins are evident in the *C. sorokiniana* genome, GTG1 and CHLH orthologs are present (Fig. 1 and 2). Additionally, *C. sorokiniana* encodes an ortholog to ABI2 (ABA insensitive 2), a protein phosphatase regulated by SOS2 that physically interacts with PYR1 in *Arabidopsis;* this ortholog is differentially regulated by ABA in *Chlorella* (Supplementary File 1). Other downstream signaling components are also present, including an ortholog to SnRK2, a stomatal closure mediating protein in higher plants (Supplementary File 1). Furthermore, *Chlorella sorokiniana* encodes the ABA biosynthetic and degradation pathway, as well as a putative ABA transporter (Table 1; Fig. 5). The genome of *C. sorokiniana* contains putative orthologs to genes in ABA mediated and non-ABA mediated stress responses in higher plants, including MKP1 and a complete SOS pathway (Table 2). *C. sorokiniana* encodes genes which are involved not only in ABA signaling, but also in signal transduction and integration with other signaling pathways.

**Table 1.** The *Chlorella sorokiniana* genome encodes orthologs to ABA biosynthesis, perception, transport, and degradation. Representative members of the respective pathways were used to conduct reciprocal BLAST searches. It was found that *C. sorokiniana* contains at least one ortholog to each pathway.

| Gene name | Function | Arabidopsis accession | *Chlorella* ortholog | Expect |
| --- | --- | --- | --- | --- |
| GTG1 | alternative ABA receptor | AT1G64990 | sca012.g110550.t1 | 6e-95 |
| ChlH | Magnesium chelatase subunit H, alternate ABA receptor, chloroplastic | AT5G13630 | sca051.g100950.t1 | 0.0 |
| NCED2 | ABA biosynthesis, 9-cis-epoxycarotenoid dioxygenase | AT4G18350 | sca015.g107700.t1 | 2e-65 |
| Abscisic acid 8'-hydroxylase | ABA inactivation enzyme | AT5G45340 | sca221.g100650.t2 | 2e-42 |
| ABCG25 | ABA efflux transporter | AT1G71960 | sca052.g100950.t3 | 1e-102 |

**Table 2.** Stress signal transduction genes in *C. sorokiniana*. The SOS pathway is found in *Chlorella*, suggesting that it is an ancient pathway that has since differentiated and specialized in higher plants. Additionally, CsMAPKKK/CsNPK1 are significantly (2.52log_2_, q = 0.027) upregulated by treatment with ABA (Table 3).

| Gene | *Arabidopsis*<br>accession | *Chlorella*<br>accession | Expect | Function |
| --- | --- | --- | --- | --- |
| SOS2 (SNRK3) | AT5G35410 | sca154.g100300.t<br>1 | 3e-119 | Ser/Thr kinase,<br>regulated by ABI2,<br>SOS3, Ca<sup>2+</sup> |
| SOS3 | AT5G24270 | sca106.g102500.t<br>1 | 3e-56 | Calcium sensor;<br>activates SOS2 |
| SOS1 (ATNHX7) | AT2G01980 | sca004.g105100.t<br>1 | 2e-80 | Na<sup>+</sup>/H<sup>+</sup> antiporter,<br>regulated by SOS2<br>and SOS3 |
| SOS4 | AT5G37850 | sca081.g103450.t<br>1 | 7e-105 | Pyridoxal kinase |
| MKP1 | AT3G55270 | sca022.g103900.t<br>1 | 8e-77 | Negative regulator<br>of salt tolerance |
| MAPKKK<br>(NPK1) | AT3G06030 | sca015.g103350.t<br>1 | 3e-109 | Activates oxidative<br>signal cascade;<br>*Arabidopsis*<br>transgenics show<br>higher<br>photosynthetic<br>rates; tolerance to<br>cold, heat, salinity |

**Figure 1.**
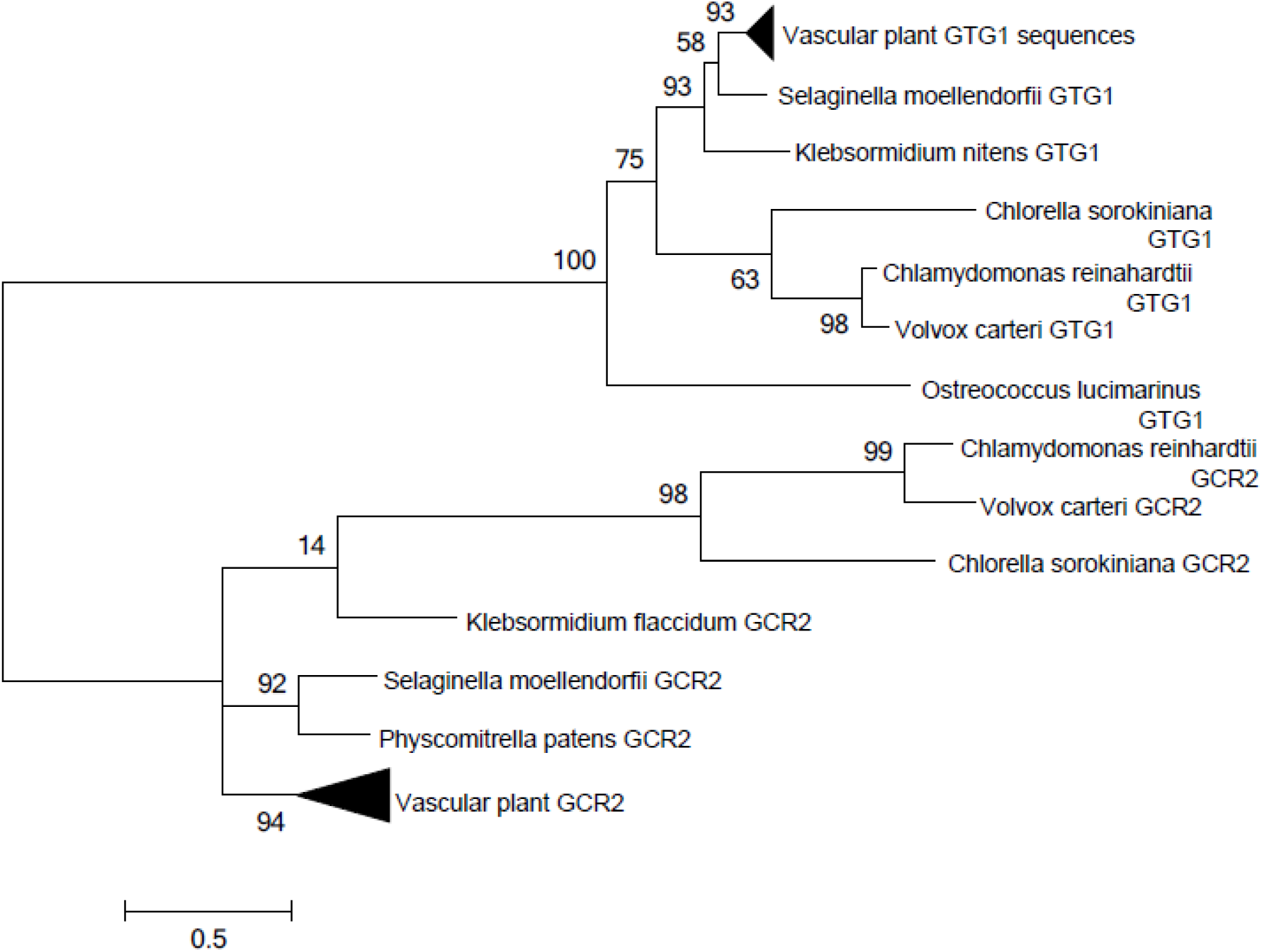
Phylogenetic analysis of GTG1, compared to GCR2, another GPCR-like protein in plants. The evolutionary history was inferred by using the Maximum Likelihood method based on the Le_Gascuel_2008 model [1]. The tree with the highest log likelihood is shown. The percentage of trees in which the associated taxa clustered together is shown next to the branches. Initial tree(s) for the heuristic search were obtained by applying the Neighbor-Joining method to a matrix of pairwise distances estimated using a JTT model. The rate variation model allowed for some sites to be evolutionarily invariable ([+I]) sites. The tree is drawn to scale, with branch lengths measured in the number of substitutions per site.

**Figure 2.**
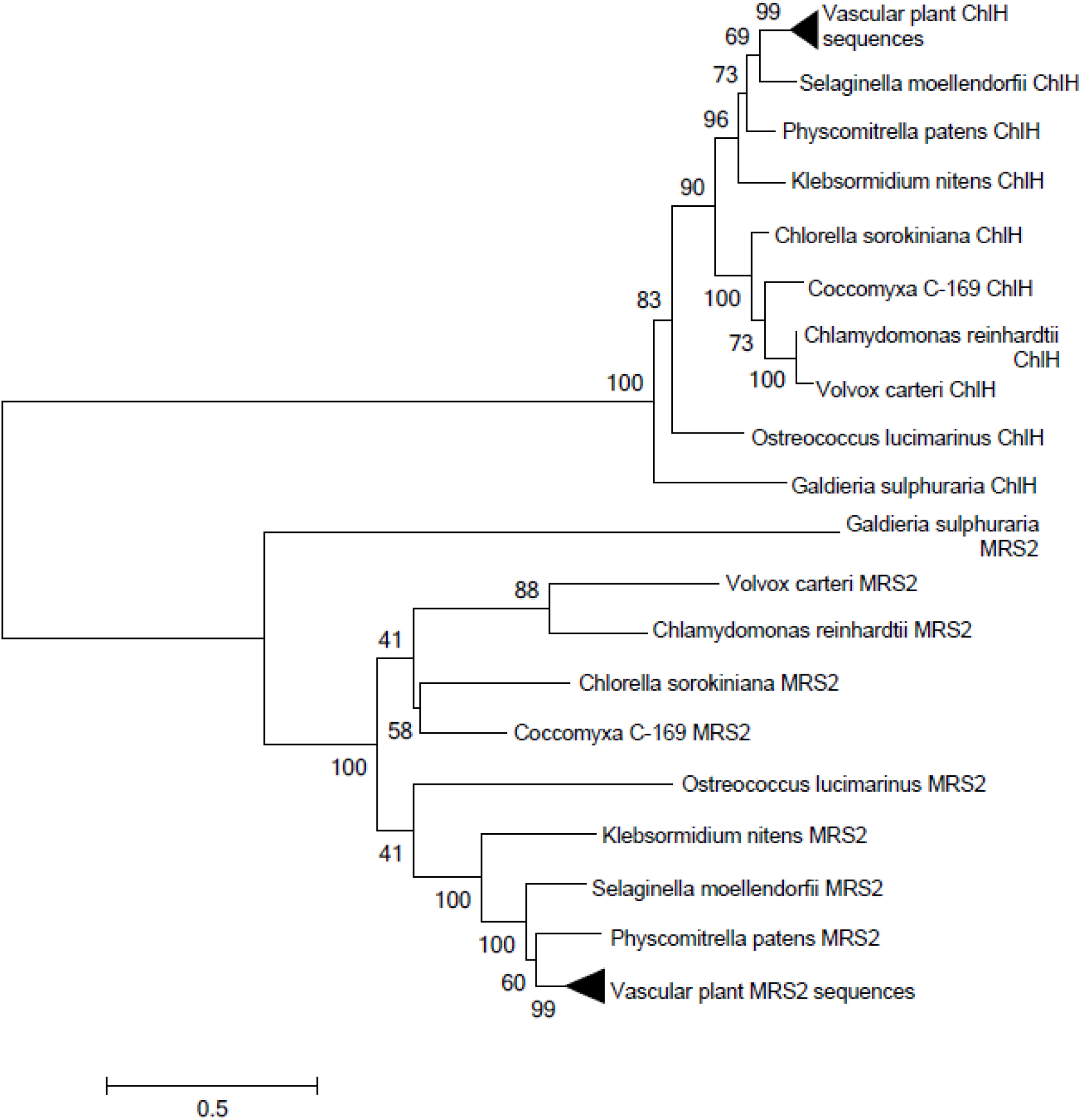
Phylogenetic analysis of ChlH, compared to MRS2, another magnesium chelatase in plants. The evolutionary history was inferred by using the Maximum Likelihood method based on the Le_Gascuel_2008 model. The tree with the highest log likelihood is shown. The percentage of trees in which the associated taxa clustered together is shown next to the branches. Initial tree(s) for the heuristic search were obtained by applying the Neighbor-Joining method to a matrix of pairwise distances estimated using a JTT model. The rate variation model allowed for some sites to be evolutionarily invariable ([+I]) sites. The tree is drawn to scale, with branch lengths measured in the number of substitutions per site.

**Figure 3.**
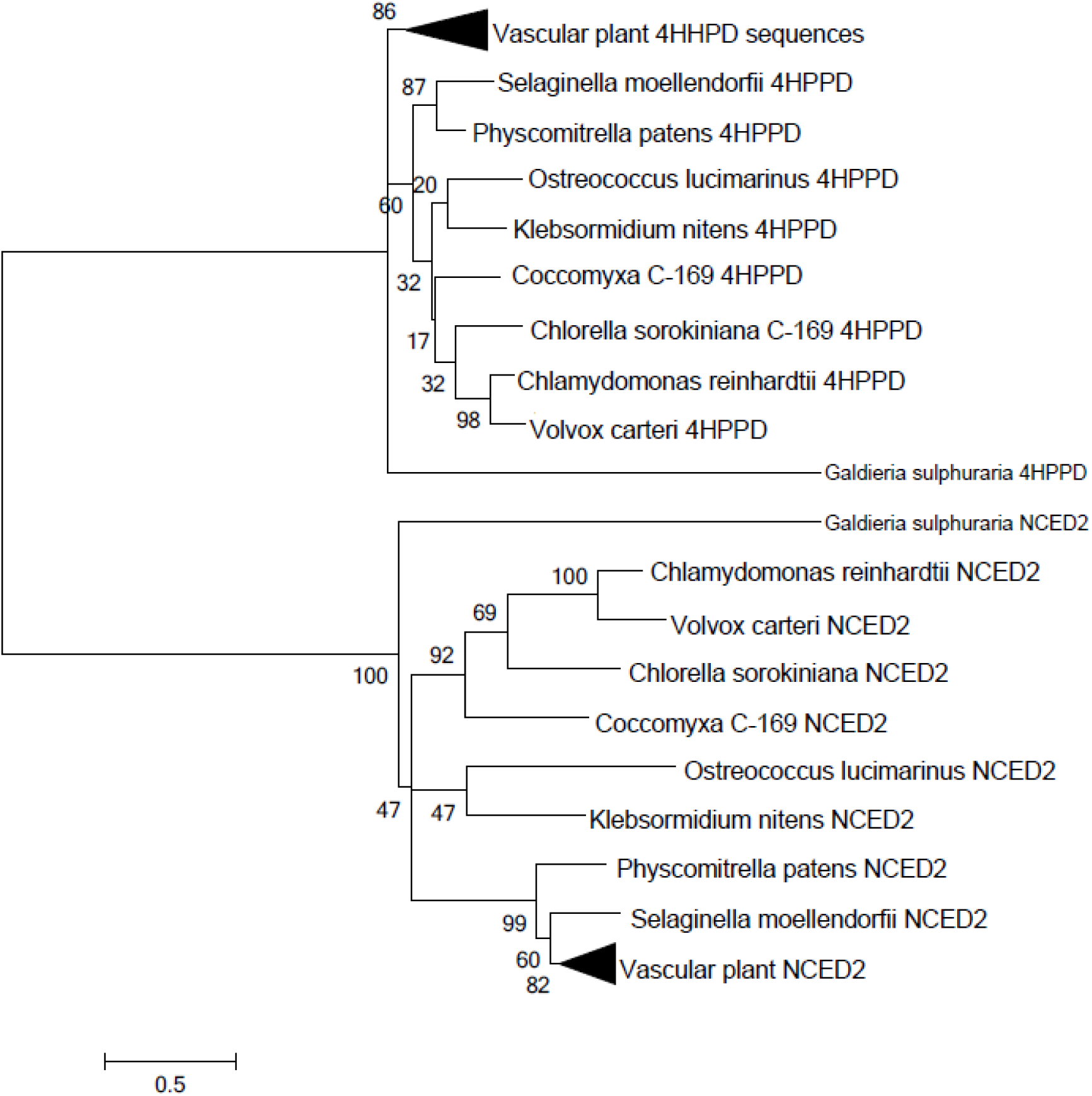
Phylogenetic analysis of NCED2, the commitment step of ABA biosynthesis, compared to 4-hydroxyphenylpyruviate dioxygenase. The evolutionary history was inferred by using the Maximum Likelihood method based on the Le_Gascuel_2008 model. The tree with the highest log likelihood is shown. The percentage of trees in which the associated taxa clustered together is shown next to the branches. Initial tree(s) for the heuristic search were obtained by applying the Neighbor-Joining method to a matrix of pairwise distances estimated using a JTT model. The rate variation model allowed for some sites to be evolutionarily invariable ([+I]) sites. The tree is drawn to scale, with branch lengths measured in the number of substitutions per site.

**Figure 4.**
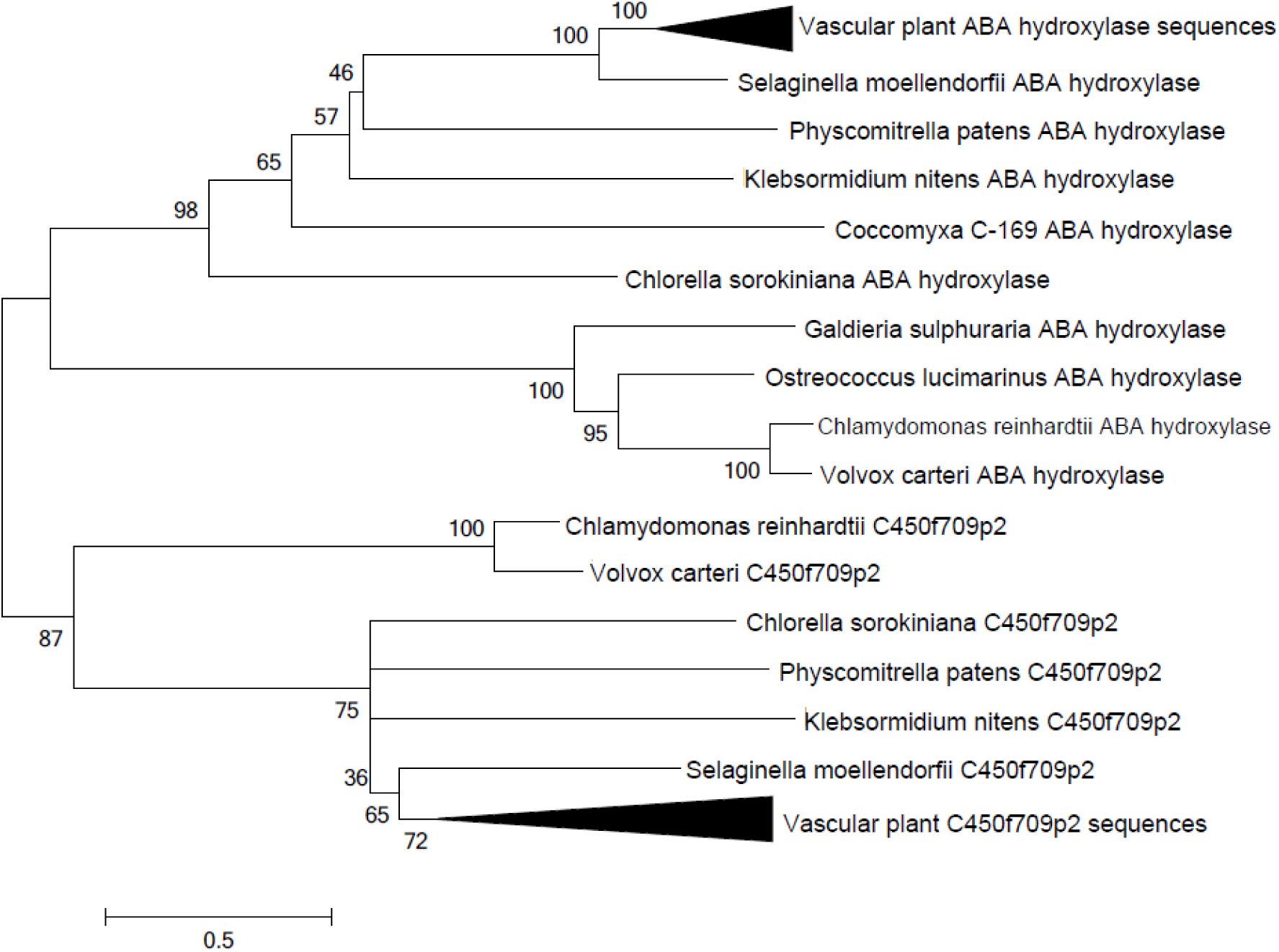
Phylogenetic analysis of ABA hydroxylase, an ABA degradation enzyme, compared to C450f709p2, a cytochrome p450 monooxygenase domain containing enzyme involved in salt tolerance. The evolutionary history was inferred by using the Maximum Likelihood method based on the Le_Gascuel_2008 model. The tree with the highest log likelihood is shown. The percentage of trees in which the associated taxa clustered together is shown next to the branches. Initial tree(s) for the heuristic search were obtained by applying the Neighbor-Joining method to a matrix of pairwise distances estimated using a JTT model. The rate variation model allowed for some sites to be evolutionarily invariable ([+I]) sites. The tree is drawn to scale, with branch lengths measured in the number of substitutions per site.

**Figure 5.**
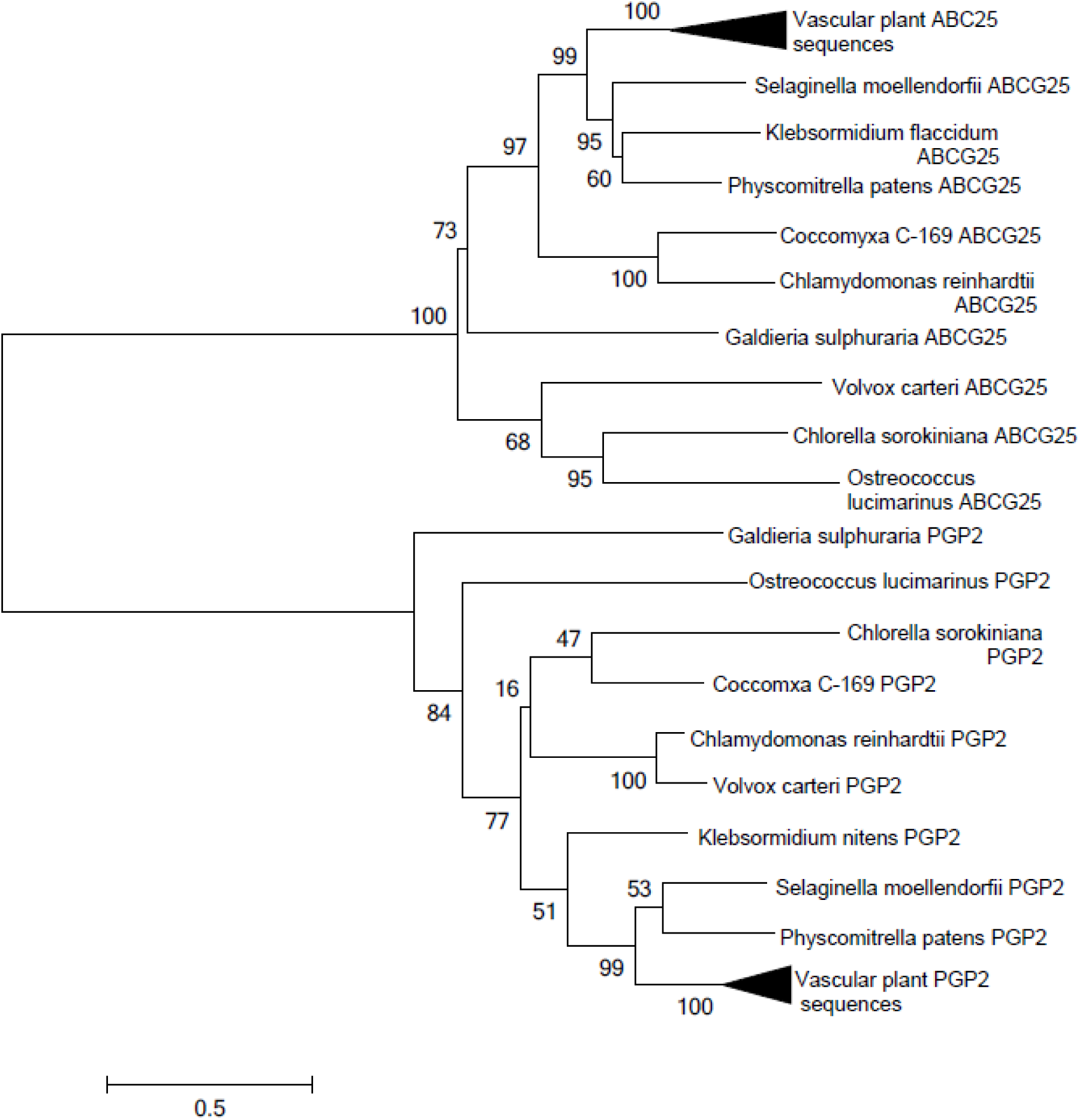
Phylogenetic analysis of ABCG25, an ABA transporter, compared to P-glycoprotein 2, a similar MDR-like transporter. The evolutionary history was inferred by using the Maximum Likelihood method based on the Le_Gascuel_2008 model. The tree with the highest log likelihood is shown. The percentage of trees in which the associated taxa clustered together is shown next to the branches. Initial tree(s) for the heuristic search were obtained by applying the Neighbor-Joining method to a matrix of pairwise distances estimated using a JTT model. The rate variation model allowed for some sites to be evolutionarily invariable ([+I]) sites. The tree is drawn to scale, with branch lengths measured in the number of substitutions per site.

### Abscisic acid and OPDA are synthesized and secreted by *Chlorella sorokiniana* during saline and oxidative stress

Plants produce ABA in response to water availability stressors, including cold, salinity, and drought. In order to determine whether *C. sorokiniana*, a freshwater alga, makes abscisic acid under similar conditions, we exposed cultures to 600 mM NaCl (the osmotic strength of seawater). Significant quantities of ABA were present in both the cell pellet and the supernatant (∼8 and 16 nM, respectively), indicating the active synthesis and secretion of ABA upon osmotic stress (Fig. 6). Additionally, the plant stress hormone OPDA was produced and secreted in significant quantities (50 nM) in the stressed cultures, but essentially absent (∼32-fold less) from the unstressed controls (Fig. 7).

**Figure 6.**
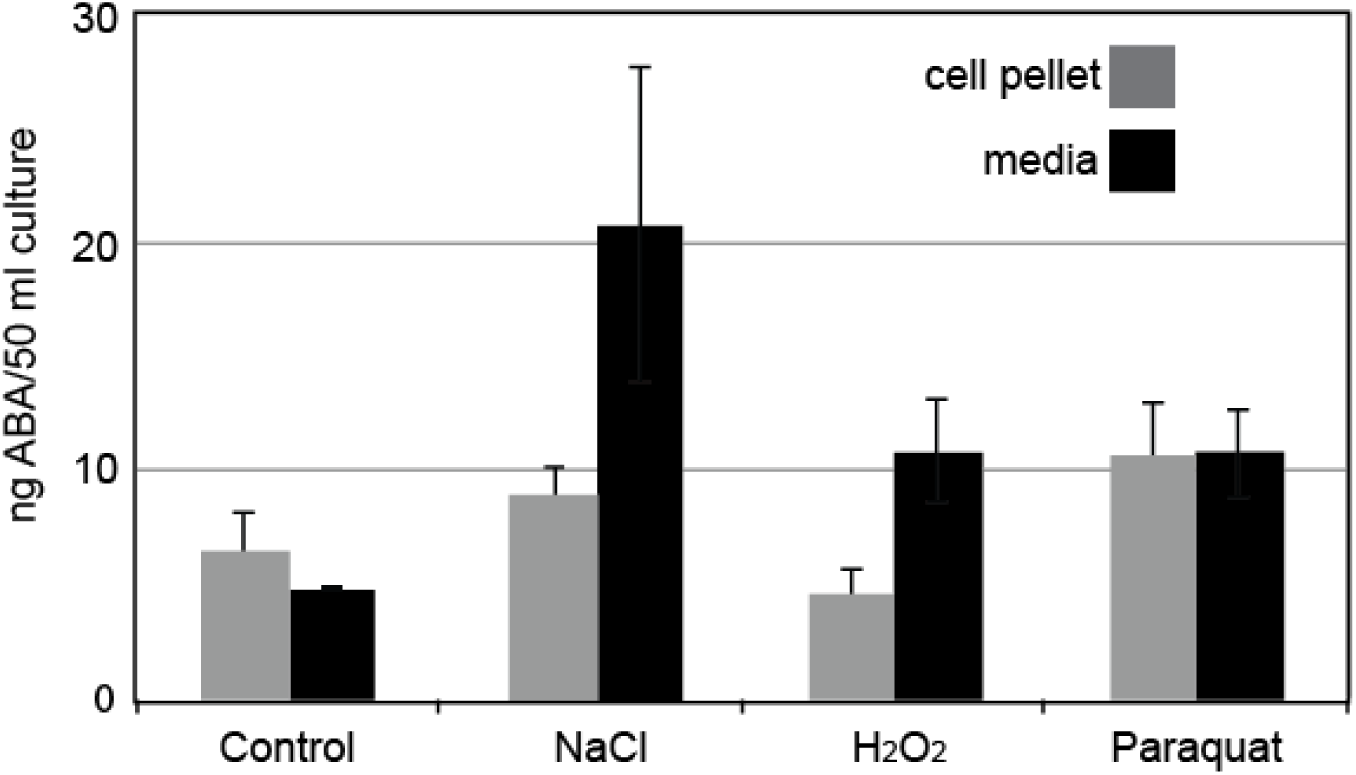
*Chlorella sorokiniana* cells produce and secrete ABA. In response to 600 mM salt stress and oxidative stresses, *Chlorella* secretes almost twice as much ABA into the supernatant as in the pellet, except for paraquat, which remains stable. Error bars represent standard error of biological replicates conducted in duplicate. Cell pellets, n =2, culture media, n = 3.

**Figure 7.**
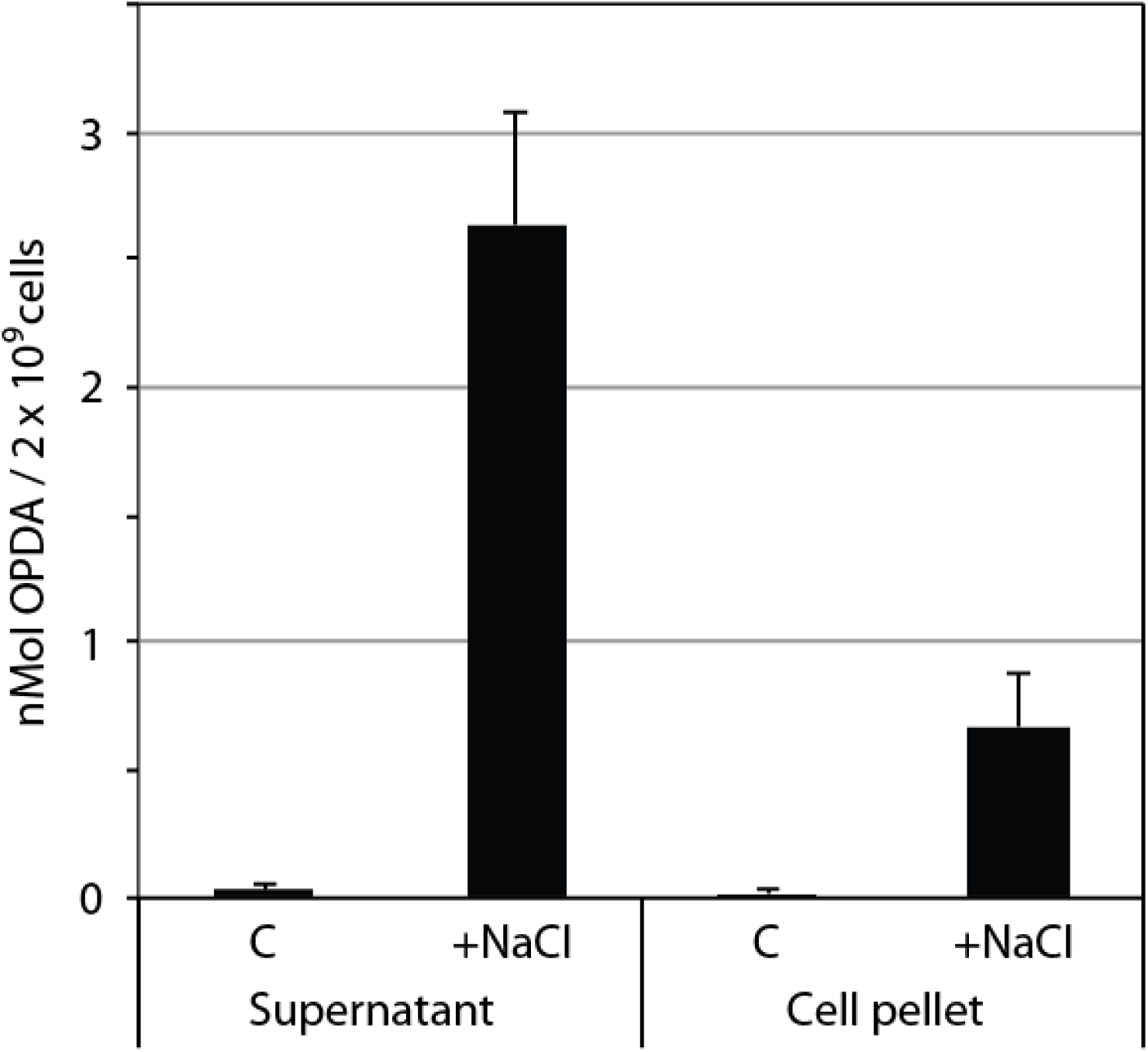
OPDA, a stress phytohormone, is produced in response to salt shock. Cultures were treated with 600 mM NaCl as described for Figure 6, and OPDA quantified by LC/MS/MS as described in Materials and Methods. Error bars represent the S.E.M. of 3 independent biological replicates conducted in duplicate.

Treatment with 1 mM H_2_O_2_ or 5 μM paraquat also increased the production of ABA in the cell pellet and supernatant (Fig. 6). In seed plants, singlet oxygen stress (stimulated by application of paraquat) directly results in the activation of ABA signaling pathways, while H_2_O_2_ signaling occurs downstream of ABA signaling and participates in the signal transduction cascade that leads to stomatal closure, as reviewed by Bright et al [25]. In *Chlorella*, both of these stressors resulted in modest induction (∼2-fold) of ABA production.

### Abscisic acid results in the differential regulation of 290 transcripts in *C. sorokiniana*

Treatment of *C. sorokiniana* cultures with 100 μM ABA for 48 h resulted in 219 upregulated and 71 downregulated transcripts. These transcripts were differentially expressed at least two-fold, with a q-value of < 0.05 as determined by CuffDiff2 [26]. A large fraction of the upregulated genes control DNA replication and repair (Fig. 8). Notably, DNA pol III, BRCA1, translesion DNA polymerase κ, and the MCM8/MCM9 orthologs participating in homologous recombination repair are upregulated as shown in Fig 9A [27]. A summary of representative genes is presented in Table 3, and a complete list of differentially regulated genes is provided as a Supplementary Data file.

Central metabolism genes upregulated by ABA.

Central metabolism genes upregulated by ABA
| Gene | *Chlorella* accession | log<sub>2</sub>-fold-change | Pathway/function |
| --- | --- | --- | --- |
| Malate synthase | sca085.g100800.t1 | 3.65 | Glyoxylate bypass |
| Pyruvate ferredoxin/flavodoxin oxidoreductase | sca098.g101900.t1 | 2.78 | Glycolysis |
| PEP carboxykinase | sca027.g100650.t1 | 2.71 | Gluconeogenesis |

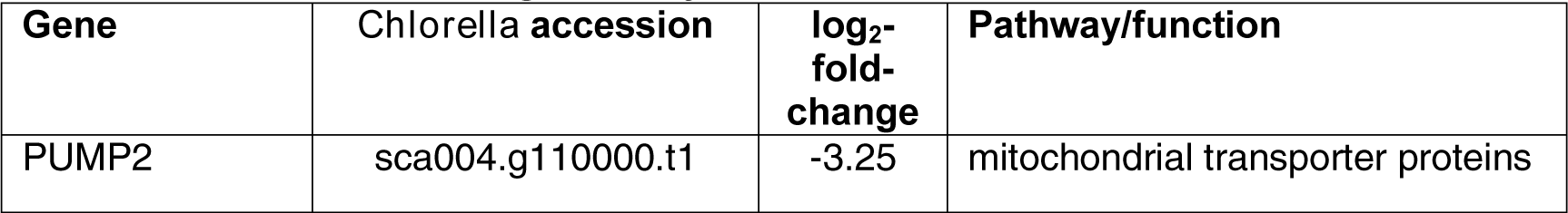
Central metabolism downregulated by ABA.

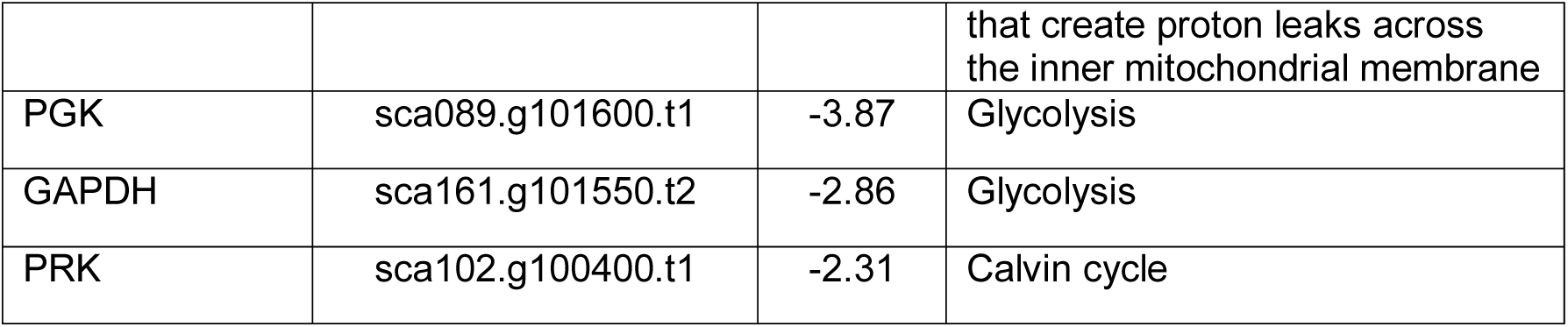
Photosynthesis genes downregulated by ABA.

**Table 3.** Representative members of gene ontology classes that are altered by ABA exposure. **DNA replication and repair proteins upregulated by ABA**

| Gene | *Chlorella* accession | log<sub>2</sub>-fold-change | Pathway/function |
| --- | --- | --- | --- |
| Carbonic anhydrase | sca004.g108450.t1 | -3.79 | Carbon concentration/photosynthesis |
| Phytoene desaturase | sca153.g100750.t1 | -3.15 | Chlorophyll biosynthesis/photosynthesis |
| Rubisco large subunit methyltransferase | sca221.g102000.t1 | -2.57 | Photosynthesis |
| Photosystem II stability/assembly factor HCF136 | sca134.g104050.t1 | -2.82 | Photosynthesis |
| Divinyl chlorophyllide A 8-vinyl-reductase | sca034.g104550.t1 | -2.38 | Photosynthesis/chlorophyll biosynthesis |
| Uroporphyrinogen-III decarboxylase | sca067.g100650.t1 | -2.3237 | Photosynthesis/chlorophyll biosynthesis |
| Coproporphyrinogen III oxidase | sca077.g103150.t1 | -2.18 | Chlorophyll/heme biosynthesis |

**Table 3 (cont.) Representative members of gene ontology classes that are altered by ABA exposure.**
| Gene | *Chlorella* accession | log<sub>2</sub>-fold-change | Pathway/function |
| --- | --- | --- | --- |
| DNA helicase MCM8 | sca081.g104500.t1 | 3.71 | **Recombination repair, with MCM9** |
| DNA helicase MCM9 | sca104.g106050.t1 | 3.37 | **Recombination repair, with MCM8** |
| DNA Pol α | sca079.g105350.t1 | 3.47 | **Okazaki fragment synthesis** |
| DNA Pol $\kappa$ | sca106.g112250.t2 | 2.48 | **DNA repair: translesion synthesis** |
| BRCA1-like | sca206.g103000.t1 | 3.69 | **Double strand break repair**, homologous recombination |

In seed plants, ABA retards growth by initiating stomatal closure via H_2_O_2_ production, thereby decreasing carbon dioxide accumulation; additionally, ABA represses photosynthesis. Accordingly, we observed a decreased abundance of transcripts associated with photosynthesis, including carbonic anhydrase, phytoene desaturase, and at least 4 chlorophyll biosynthesis associated genes including magnesium chelatase subunit ChlH [12] (Table 3). ChlH also acts as a chloroplastic ABA receptor in some plants: in *Arabidopsis*, it specifically binds ABA and regulates a cascade of ABA signaling genes including RD29A, MYB2, and MYC2 among others [28]. It is important to note that, unlike in higher plants, we observed no decrease in cell number or biomass yield in response to ABA treatment under our growth conditions.

Central metabolism enzymes, including PGK, GAPDH, PRK, and malate synthase, were downregulated by ABA. In *Arabidopsis*, mutant lines with specific defects in glycolysis have altered ABA signal transduction: *Arabidopsis* plants deficient in plastidial glyceraldehyde-3-phosphate dehydrogenase (GAPCp) are ABA insensitive in stomatal closure, growth, and germination assays [29]. Indeed, the downregulation in the lower portion of the Embden-Meyerhof-Parnas pathway may suggest the shunting of carbon towards glycerol, a common compatible osmolyte in algae. Muñoz-Bertomeu et al. demonstrate that the disruption of sugar and serine homeostasis causes ABA insensitivity in *gapcp1gapcp2* plants, and supplementing sugar and serine restored and even augmented ABA sensitivity (18).

In addition to direct effects on carbon fluxes and energy capture, altered expression of glycolysis and photosynthesis related genes have been shown to alter the production of ROS, suggesting that ROS signaling is integrated with primary cellular metabolism[15,30]. Crosstalk between ROS and ABA signaling has been demonstrated in plants, where it is influenced by altered metabolic fluxes. Our results sugest that ROS production and scavenging are also altered, with coordinate effects on ABA signaling [20] as previously reported [15].

Given that a major stress response of many algae is the accumulation and storage of fatty acids in lipid droplets [31], it was surprising that fatty acid biosynthetic genes were largely downregulated by ABA. These genes, including 3-oxoacyl ACP synthase, acyl carrier protein, biotin carboxyl carrier protein, and biotin carboxylase. These components contribute to both fatty acid synthase and acetyl-CoA carboxylase functions in chloroplasts, and suggest the likelihood of additional alterations in central metabolic pathways upon treatment with ABA. GPI (glycero-phospho-myo-inositol) ethanolamine phosphate transferase 3, participating in GPI anchor biosynthesis, was one of very few lipid-active genes to be upregulated by ABA, perhaps indicating an alteration in the abundance or type of GPI-anchored proteins displayed at the cell surface. Overall, carbon flow appears to be shunted away from lipid biosynthesis, at least as judged by steady-state levels of associated transcripts.

Several putative ABA signal transduction genes are differentially expressed in *Chlorella* upon exposure to ABA. These include ChlH, a chloroplastic ABA receptor repressed by ABA. Additionally, an ortholog to MAP3K/NPK1, a kinase which in *Arabidopsis* regulates the drought stress response downstream of H_2_O_2_ signaling, is upregulated by ABA (Supplementary file 1). Notably, the *Arabidopsis* NPK1 ortholog ANP1 induces MPK3 and MPK6 expression via H_2_O_2_, so that the MPK3 and MPK6 can then participate in genotoxic and drought stress tolerance. In *Arabidopsis*, MKP1 (MAP kinase phosphatase 1) also interacts with both MPK3 and MPK6 to mediate salt tolerance. In *Arabidopsis*, MKP1 is postulated to be post translationally regulated [20], and the same may be true for *Chlorella*, as differential regulation was not detected in our studies.

### ABA pretreatment induces UV tolerance

Due to the upregulation of DNA replication and repair mechanisms (Figs. 8 and 9A), we suspected that ABA may have a role in priming cells for DNA repair. We investigated this possibility by inducing DNA damage in cells with short-wave UV irradiation (18 mJ, 270 nm). For untreated cells, this dose reduced the colony forming units (i.e. viable cells) to roughly 8% of the untreated control, representing 92% killing. In contrast, the survival of cells pretreated with 10 μM ABA nearly doubled over the untreated control, to ∼15% (Fig. 9B). This result directly demonstrates that treatment with exogenous ABA primes *Chlorella* cells for DNA damage, and that the mechanism by which it does so is very likely to be the via the induction of DNA damage response genes as shown in our transcriptome analysis (Fig. 8).

**Figure 8.**
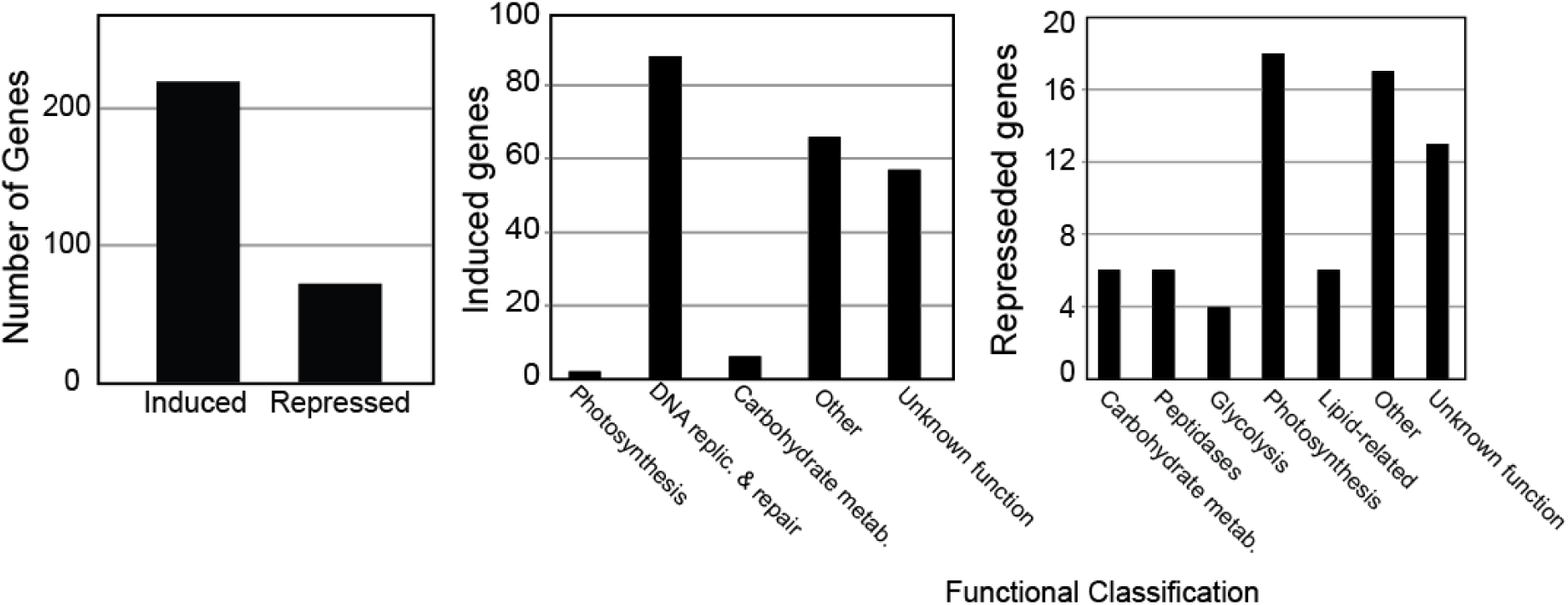
Differential expression of genes upon ABA treatment. A: 290 total genes are differentially regulated by at least twofold: 71 are downregulated, and 219 are upregulated. B. DNA replication and repair related genes make up the largest category of upregulated genes with predicted functions. C. Genes related to photosynthesis and glycolysis are downregulated.

**Figure 9.**
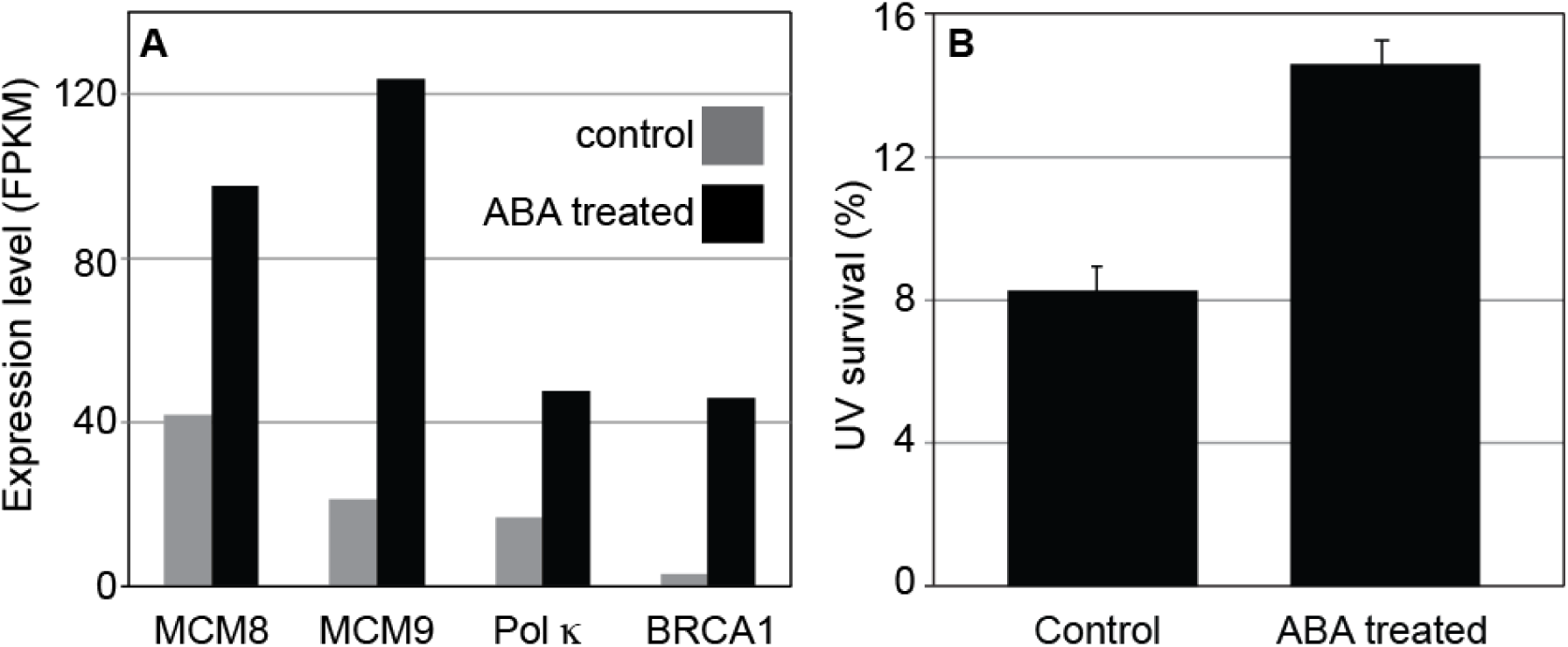
Abscisic acid pretreatment enhances tolerance to UV irradiation. A. DNA repair genes are upregulated by treatment with ABA. B. Cells pretreated for 48 hours with 10 μM ABA were found to have a survival rate of 15%, compared to an 8.3% survival rate for the untreated cells. N = 4, *p*= 0.0004, computed using the student’s T test for equal variances.

## Discussion

This work demonstrates that ABA is produced, secreted, and sensed by *Chlorella sorokiniana*, as summarized by our model in Fig. 10. Though plant hormone signaling has been proposed for some green algae based on reports of its production and presence of related gene orthologs, our study is the first to integrate genomic, transcriptomic, biochemical, and physiological evidence of algal cell to cell communication using a phytohormone. In higher plants, exposure to exogenous ABA causes plants to reduce photosynthesis and respiration. Conversely, external effects on ABA in algae have been reported to be minimal [8], consistent with our observations: *Chlorella* cells treated with up to 100 μM ABA did not experience a reduction in growth rate or cell yield.

**Figure 10.**
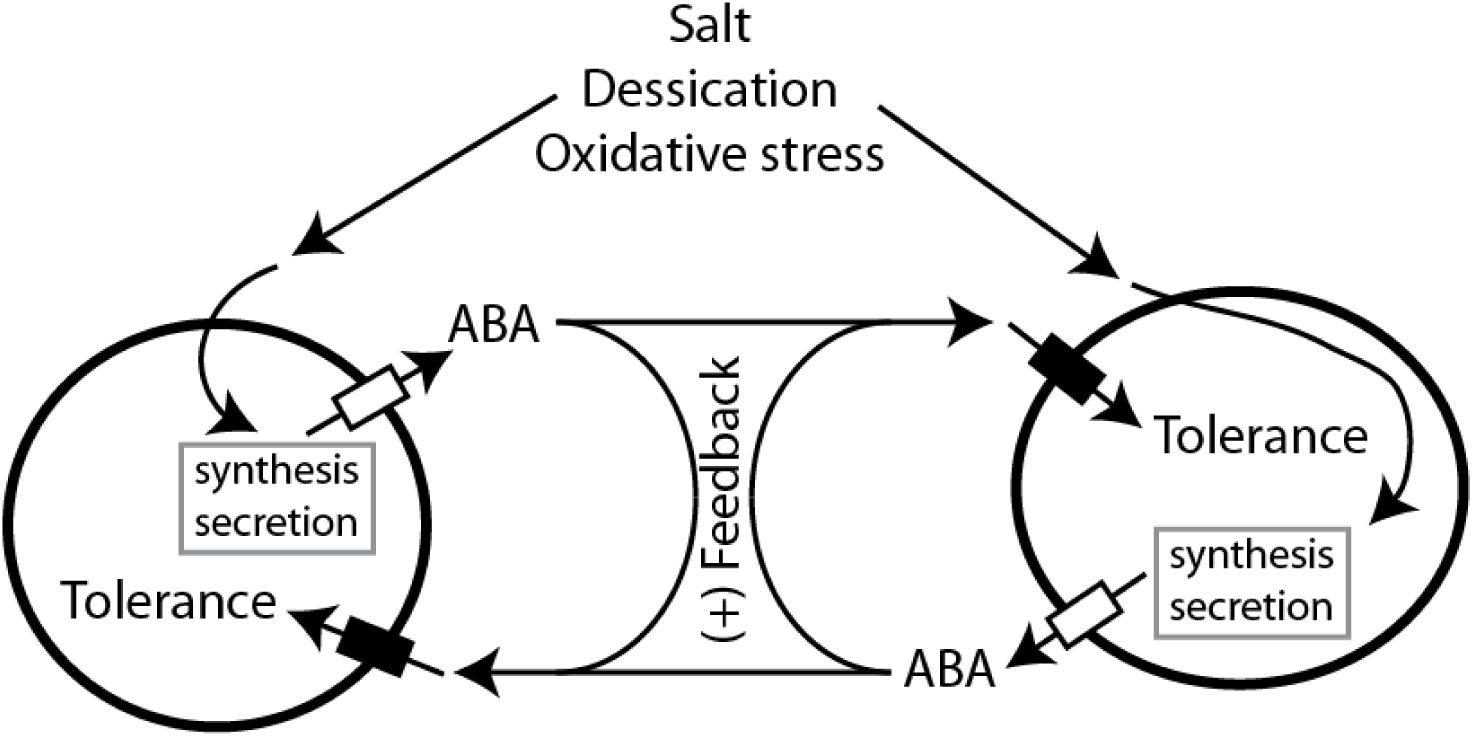
A model for ABA synthesis, secretion, and signaling in *C. sorokiniana*. We propose a framework in which abiotic stressors induce the synthesis and secretion of ABA and other potential signaling molecules, leading to gene expression changes associated with stress tolerance. Importantly, the concentration of these molecules will be proportional to the population density, allowing for signal amplification and coordination of population-level stress responses.

Phylogenetic analysis (Figs. 1-5) demonstrates that the genes listed in Tables 1-4 are orthologous to ABA synthesis and perception genes in plants, indicating that the development of ABA signaling may have been essential for the colonization of land. ABA also provides a mechanism for plants and algae to cope with the effects of salt and desiccation stress, both of which are amplified on land. Phytohormones have been implicated in the desiccation tolerance of *Klebsormidium* species, which are filamentous charophyte algae that are thought of as a bridge between unicellular and multicellular plants. ABA related genes in *K. flacidum* were upregulated in response to desiccation, though the hormone itself was not quantified in the study [32]. Holzinger and Becker noted, as we do, that our respective organisms contain almost complete ABA signaling pathways except for the PYR/PYL/RCAR type receptors, AREB family transcription factors, and S-type anion channel SLAC1 that is required for stomatal function in higher plants [33]. Transcriptomic studies in *Klebsormidium* have revealed a land plant-like defense reaction against desiccation. Indeed, the presence of certain components of these pathways suggests that the use of ABA as a stress signaling molecule by plants occurred early in the evolutionary history of plants – at least as early as the unicellular ancestors of charophytes and chlorophytes – and that it is regulated by ancient gene families which have since differentiated and specify multiple stress and developmental functions in higher plants (32, 34).

Components of ABA signaling, including stomatal closure regulating protein SnRK2 are present in *Chlorella*, and appear to be differentially expressed upon treatment with ABA. *Chlorella* also contains a putative ortholog to MAP kinase phosphatase 1 (MKP1), which participates in salt stress adaptation and tolerance in *Arabidopsis* [20]. It is possible that it is regulated at an earlier stage after ABA application than what this study measured, and that early ABA signal transduction pathways were not within the scope of this study’s methodologies.

Previous studies have demonstrated that genotoxic stresses are linked to salinity and ABA signaling in *Arabidopsis* [35]. Saha et al. [36] suggested that the ROS accumulation under NaCl stress contributed to an increase in DNA damage, interconnecting the two stressors with ABA signaling We propose that, in Chlorella, salt stress triggers a signaling cascade that activates ABA, resulting in protection from DNA damage, whereas UV-induced DNA damage itself does not trigger ABA production (Fig. 10). The production of ABA in response to oxidative stress bolsters this model: salt and oxidative stresses both activate ABA signaling, possibly in addition to ABA-independent oxidative stress response and salt stress response pathways. In contrast to seed plants, both paraquat and hydrogen peroxide stresses caused ABA accumulation, suggesting altered mechanisms of cross-pathway regulation between oxidative stress and salt signaling pathways in this alga relative to seed plants. This parallels studies in auxin signaling in algae, in which canonical transcriptional auxin receptors (TIR1/AFB) are absent, but alternate receptors are present (ABP1) (37, 38).

As the interrelation of abiotic stressors and signaling pathways is an emerging field in plant stress biology, we propose that *Chlorella* will provide a convenient unicellular model system for elucidating the interrelationship of salt and oxidative stresses, DNA damage, and ABA signaling in plant cells (Fig 10). This study also demonstrates that ABA signaling can occur without PYR/PYL/RCAR signaling as it is currently understood, and we propose that the study of an organism with a well-developed ABA response, but missing PYR/PYL/RCAR, could offer advantages to the study of these ancestral modes of ABA perception and signaling, and will help foster a better understanding of ABA signaling in a functional and evolutionary context. In this vein, we note that *Chlorella* encodes an ortholog to ABI2 (ABA insensitive 2), a protein phosphatase regulated by SOS2 that physically interacts with PYR1 in *Arabidopsis* (Park 2009); this ABI2 ortholog is differentially regulated by ABA in *Chlorella* (Table 2), suggesting that PYR/PYL/RCAR-related components are present, but that their functions were recruited for ABA signaling at a later stage in the evolution of higher plants. This observation suggests further interactions between the SOS and ABA pathways, and provides a putative connection between ABA and SOS signaling and the evolution of the PYR/PYL/RCAR family of receptors.

In summary, we demonstrate that ABA has a physiological role in the chlorophyte alga *Chlorella sorokiniana*, and we demonstrate its role as an intercellular stress signaling molecule in this organism. Exogenous ABA primes cells to resist DNA damage by upregulating DNA replication and repair related transcripts. The use of ABA as a signaling molecule and the interrelation of drought and UV stress suggest that ABA was a critical step in the evolutionary advancement of land plants. These findings fit well with other current work from our laboratory [38] on auxin (indole-3- acetic acid) signaling in unicellular chlorophytes. Specifically, we have identified a suite of genes in *C. sorokiniana* that facilitate the synthesis, secretion, and perception of auxin by this organism. These findings, coupled to our current studies on ABA signaling, provide strong evidence that multiple phytohormone signaling pathways were present and operative in unicellular algae prior to the evolution of multicellularity and colonization of terrestrial environments, and that these pathways were expanded and diversified during the subsequent evolution of seed plants.

## Materials and Methods

### Culture conditions

An agar slant of *Chlorella sorokiniana* UTEX 1230 culture was obtained from the University of Texas culture collection on proteose peptone media. Long term cultures were maintained on TAP agar plates. Starter cultures for all experiments were allowed to grow to saturation in Bold’s Basal Media (BBM) prior to dilution for commencement of experiments. All experiments were conducted with at least three biological replicates; each biological replicate contained two technical replicates.

### Salt Stress

A saturated starter culture was used to inoculate four flasks per experiment (two each of a control and a salt stressed flask). 250-mL flasks containing 50 mL of culture at 5×10^6^ cells/mL were prepared for salt stress. After 48 hours of growth (approximately 2.5 doublings), with an average cell density of 3-4×10^7^ cells/mL, cells to be treated were centrifuged at 2,500 x *g* for five minutes and resuspended in either fresh BBM, or BBM containing 600 mM NaCl. Cells were harvested after 16 hours of salt treatment and processed immediately for RNA isolation or compound quantification.

### Oxidative stress

A saturated starter culture was used to start four flasks per experiment (two each of a control and a salt stressed flask). 250-mL flasks containing 50 mL of culture at 5×10^6^ cells/mL were set up for salt stress. After 48 hours of growth (approximately 2.5 doublings), with an average cell density of 3-4×10^7^ cells/mL, 5 μM paraquat (from a 5 mM stock in water) and 1 mM H_2_O_2_ (from a 1M stock) were added. Cells and supernatants were harvested after 16 hours of oxidative stress treatment.

### Hormone extraction and quantification

*ABA recovery from spent medium.* Culture supernatants were harvested by centrifugation and vacuum filtered through a 0.22 μm filter. 1 ng/ml of deuterated ABA internal standard (d_6_-ABA; [^2^H_6_](+)-cis,trans-abscisic acid, OlChemIM, Olomouc, Czech Republic) was added to each sample. The supernatant was passed through a Supelco Discovery C18 reverse phase column (Cat. no. 52604-U) according to the manufacturer’s protocol, and eluted in a buffer of 50% acetonitrile/1% acetic acid. Upon elution, samples were immediately stored at −80°C until vacuum concentration and analysis by LC/MS/MS.

*Extraction of cell pellets.* Pellets of 1-2 × 10^9^ cells were extracted by agitation in 10 mL of an 80% acetonitrile/1% acetic acid solvent overnight at 4°C. After centrifugation to remove cell debris, the extract was evaporated under N_2_ gas in an N-Evap^TM^ 112 (Organomation Associates, Inc.) and resuspended in ∼20 mL of water. This preparation was subsequently passed over a C18 reverse phase column and eluted as described above.

*LC/MS/MS quantitation parameters.* For LC separation, a ZORBAX Eclipse Plus C18 reversed-phase column (2.1 mm × 100 mm, Agilent) was used flowing at 0.45 mL/min at 40°C. The LC system is interfaced with a Sciex QTRAP 6500+ mass spectrometer equipped with a TurboIonSpray (TIS) electrospray ion source. The hormones ABA, SA, JA, JA-Ile, and OPDA were analyzed in negative ion mode using the following source parameters: curtain gas, 20 arbitrary units (a.u.); source gas 1, 50 a.u.; source gas 2, 50 a.u.; collision activated dissociation, high; interface heater, on; temperature, 500°C; ionspray voltage, –4500. The LC gradient was from 80% solvent A (0.1% [v/v] acetic acid in Milli-Q water) to 50% A in 0.5 min, then to 100% solvent B (90% acetonitrile [v/v] with 0.1% acetic acid [v/v]) in 3.5 min, then hold for another 3.5 min at 100% B. The LC was finally ramped back to initial conditions in 0.5 min and re-equilibrated for both methods for 30 min. Both quadrupoles (Q1 and Q3) were set to unit resolution. Analyst software (version 1.6.3) was used to control sample acquisition and data analysis. The QTRAP 6500+ mass spectrometer was tuned and calibrated according to the manufacturer’s recommendations. All hormones were detected using MRM transitions that were previously optimized using a standard and a deuterium-labeled standard. For quantification, a series of standard samples containing different concentrations of unlabeled hormones was prepared. The peak area in samples was first normalized in the same way as used for the standard samples and then quantified according to the standard curve.

*OPDA quantitation.* Supernatants were harvested as described above, then evaporated under reduced pressure with gentle heating in a rotary evaporator (Rotovapor R-215, Büchi). In order to recover secreted metabolites, 2 ml methanol was then swirled around the flask and transferred to a 1.5 mL conical tube. The tube was centrifuged to remove any insoluble carryover, and stored at −20°C until LC/MS/MS analysis as described in [39].

### UV Irradiation

A saturated starter culture was diluted to 5×10^6^ cells/mL. Two flasks were set up for treatment with 10 μM ABA dissolved in methanol from a 10 mM stock at a 1:1000 dilution. The control included methanol at a 1:1000 dilution. Methanol was not found to affect any growth properties at this concentration. After two days of growth, at an average cell density of 3-4*10^7^ cells/mL, cells were diluted to 1*10^6^ cells/mL, and 10 mL of this low density cell mixture was transferred to 5 mL sterile petri dishes, resulting in a thin layer that would avoid shading from subsequent UV exposure. Cells were exposed to UV treatment in a UV Stratalinker (model, part number)

### Phylogenetic analyses

Phylogenetic trees were constructed using either TCoffee [40] or Clustal-Omega [41]. Maximum likelihood trees were generated and visualized using the LG + I model with Mega 6.06[42]. The orthologs of each gene were identified by BLAST searches of NCBI and other genome-specific databases. Outgroups for rooting of trees were selected based on similarity of primary sequence and similar enzymatic or transport functions.

### Transcriptomic analysis

A 50 mL culture of *C. sorokiniana* in BBM was grown to saturation and used to inoculate experimental cultures at a density of 5 × 10^6^ cells/ml in 125 mL media in a 1L flask. Cells were grown for 48 hours in continuous light to a density of 3-4 × 10^7^, with 100 μM ABA (added from a 100x stock in methanol) and with an ABA-free methanol control, in duplicate. Cells from 100 ml of culture were harvested by centrifugation, frozen in liquid nitrogen, and stored at −80°C until isolation. RNA was isolated using TriZOL (Invitrogen) as described in the manufacturer’s protocol. Library preparation and sequencing (50 basepair single-end reads) was carried out by Cofactor Genomics (St. Louis, MO) on the Illumina Genome Analyzer 2000 platform.

We applied a stringent quality filter process because sequencing errors can cause difficulties for the assembly algorithm [43]. The Illumina single-end reads that did not have a minimum quality score of 20 per base (Q20, corresponding to a 1% expected error rate) across the whole read were removed using PRINSEQ [44], and any unknown nucleotides (N) were trimmed. Total read counts after removal of low-quality sequences, per sample, were: Control 1: 37,320,193; Control 2: 47,652,886 ; ABA 1: 60,393,257; ABA 2: 53,749,656. The Illumina reads were deposited into the NCBI (National Center for Biotechnology and Information) Sequence Read Archive (SRA, http://www.ncbi.nlm.nih.gov/sra).

### Differential expression analysis

To assess differential gene expression induced by ABA treatment, the reads were mapped onto the *C. sorokiniana* genome using bowtie2 (ver. 2.2.9) [45]. Using uniquely mapped reads, cuffdiff implemented in the Cufflinks package (ver. 2.2.1) was used to calculate the relative differential expression and analyze the significances of observed changes between two groups [46]. Among 13,906 genes, 246 genes were differentially expressed (187 up-regulated and 63 down-regulated; *q*-values ≤ 0.05, False Discovery Rate < 0.05) (Supplementary file 1).

